# A remarkable expansion of oligopeptide transporter genes in rust fungi (Pucciniales) suggests a specialization in nutrients acquisition for obligate biotrophy

**DOI:** 10.1101/2022.04.20.488971

**Authors:** Pamela Guerillot, Asaf Salamov, Clémentine Louet, Emmanuelle Morin, Pascal Frey, Igor V. Grigoriev, Sébastien Duplessis

## Abstract

Nutrients acquisition by rust fungi during their biotrophic growth has been assigned to a few transporters expressed in haustorial infection structures. We performed a comparative genomic analysis of all transporter genes (hereafter termed transportome) classified according to the Transporter Classification Database (TCDB) focusing specifically on rust fungi (order Pucciniales) versus other species in the Dikarya. We also surveyed expression of transporter genes in the poplar rust fungus for which transcriptomics data are available across the whole life cycle. Despite a significant increase in gene number, rust fungi presented a reduced transportome compared to a vast majority of fungi in the Dikarya. However, a few transporter families in the subclass Porters showed significant expansions. Noteworthy, three metal transport-related families involved in the import, export and sequestration of metals were expanded in Pucciniales and expressed at various stages of the rust life cycle suggesting a tight regulation of metal homeostasis. The most remarkable gene expansion in the Pucciniales was observed for the oligopeptide transporter (OPT) family with 25 genes in average compared to seven to 14 genes in the other surveyed taxonomical ranks. A phylogenetic analysis showed several specific expansion events at the root of the order Pucciniales with subsequent expansions in rust taxonomical families. The OPT genes showed dynamic expression patterns along the rust life cycle and more particularly during infection of the poplar host tree, suggesting a specialization for the acquisition of nitrogen and sulfur through the transport of oligopeptides from the host during biotrophic growth.

## Introduction

Fungi of the order Pucciniales are plant pathogens responsible for rust diseases and are known to cause severe damages to major crops (e.g. wheat or soybean); as well as to trees used for wood production (e.g. eucalypts, pines or poplars) (Figueroa et al. 2018; Lorrain et al. 2019). Genes underlying virulence in rust fungi are widely studied in order to identify new means of resistance for deployment in Agriculture (Figueroa et al. 2016), however, other aspects of their biology are still scarcely studied at the molecular level (Duplessis et al. 2021). As obligate biotrophs, fungi of the order Pucciniales depend on their host plants for their growth and dissemination (Voegele et al. 2009). They display a unique life cycle which is the most complex known in fungi with the production of up to five different types of spores, the capacity to infect a single or two different host plants across the year and with a dormancy stage during winter under temperate climate (Duplessis et al. 2021). Rust fungi develop specific infection structures called haustoria within the cell cavities of infected plant cells. Haustoria allow both to deliver virulence effectors to interfere with the host immune system and to establish in the host; and to derive nutrients to ensure biotrophic growth (Garnica et al. 2013; Lorrain et al. 2019). They deploy adapted transport systems to acquire nutrients from the host, possibly with some level of specificity on their different host plants (Lorrain et al. 2019; Petre et al. 2020; Struck 2015; Voegele and Mendgen 2011).

Transporters allow the circulation of molecules across the permeability barrier that is the cellular membrane. They have been widely studied in eukaryotes and particularly in fungi for their specific involvement in the colonization of a large diversity of environments; for agronomical, ecological or industrial purposes; as for their role in different types of interaction with symbiotic partners (Courty and Wipf 2016; Garcia et al. 2016; Saier 1999). The transporter classification database (TCDB) categorizes the variety of conserved or unique transport systems in prokaryotes and eukaryotes (Saier et al. 2006). The TCDB is regularly updated and provides a solid support for the survey of transporters in genomic studies (Saier et al. 2021). The organization of transporters into classes and families allows a better understanding of their evolution and specialization in various organisms. How transporters localize and function at the interface between organisms has been a central theme in plant-microbe interactions and it was particularly well studied in model, mycorrhizal or pathogenic fungi (Brohée et al. 2010; Garcia et al. 2016; Perlin et al. 2014).

Only a handful of transporters have been studied in rust fungi, most of which were specifically detected or preferentially expressed in haustoria, and most often were localized at the haustorial membrane facing the extra-haustorial matrix and the host plasmalemma (Lorrain et al. 2019). *Uromyces fabae* has been a major model to study the localization, expression and substrate affinity of transporters in rust fungi (Mendgen et al. 2000; Voegele et al. 2009). The amino acid transporters AAT1, AAT2 and AAT3 (TCDB family 2.A.3; Hahn et al. 1997; Struck et al. 2002, 2004); the hexose transporter HXT1 (TCDB subfamily 2.A.1.1; Voegele et al. 2001); and the proton pump H+-ATPase PMA1 (TCDB subfamily 3.A.3.3; Struck et al. 1998) were particularly studied in the bean rust fungus. Recently, limited efforts have been deployed to functionally characterize rust fungal transporters. A homolog of the hexose transporter HXT1 was reported in the wheat rust fungus *Puccinia striiformis* f. sp. *tritici*, in which its silencing decreases fungal growth during plant colonization and reduces virulence (Chang et al. 2020). Different transcriptomic studies have highlighted the specific expression of transporters in haustoria, in germinating spores or in infected plant tissues (Duplessis et al. 2011a, b; Fernandez et al. 2012; Garnica et al. 2013; Talhinhas et al. 2014; Tremblay et al. 2013). Beside amino acid and hexose transporters, oligopeptide transporters (OPTs) were similarly reported as highly expressed in purified haustoria, suggesting that this type of transporters may represent an additional mean of gathering nutrients from the host in rust fungi (Garnica et al. 2013).

A dozen of rust fungi genomes are available for comparative genomics to study specific gene families and their role in the evolution of biotrophy in rust fungi (Aime et al. 2017; Bakkeren and Szabo 2020). The Joint Genome Institute (JGI) MycoCosm portal collects genomes across the Kingdom Fungi and proposes expert annotations allowing for broad comparisons (Grigoriev et al. 2014). The annotation of transporter genes has been implemented in MycoCosm according to the TCDB. Transporter families were previously described in pioneer rust genomic studies (i.e. the poplar rust and the wheat stem rust fungi; Duplessis et al. 2011a), but since, no systematic survey has been reported for rust fungi. Beyond genomes, complementary transcriptomic studies allow to pinpoint the transporter genes which are specifically expressed during colonization, biotrophic growth or sporulation (Duplessis et al. 2014, 2021). The transcriptome of the model rust fungus *Melampsora larici-populina* has been studied extensively along its life cycle which allows for a comprehensive survey of specific gene categories through transcriptomics of the whole life cycle (Guerillot et al. 2021).

In this study, we performed a survey of the transportome (i.e., a full collection of transporters in a given genome) in the order Pucciniales compared with other fungi in the Dikarya. We showed that the genomes of Pucciniales are marked by a global contraction of transporter gene families compared with other fungi and a few specific expansions. Strikingly, the subclass of porters which contains the previously described hexose and amino-acid transporter families, were significantly contracted. We showed that a few families exhibited significant expansions in Pucciniales compared to other fungi, among which the OPT family was the most remarkable. The systematic survey of transcriptomes available along the life cycle of the model rust fungus *M. larici-populina* reveals specific expression profiles of transport systems during host infection. These unique features might reveal a specialization in the success of the obligate biotrophic mode of these fungal pathogens.

## Material & Methods

### Genomic resources

Fungal genome resources were retrieved from the DoE Joint Genome Institute MycoCosm website as of the 2022/02/05 (https://mycocosm.jgi.doe.gov/; published genomes available for comparative studies were surveyed; Supplementary Table S1). The transporter annotation tables containing classification and annotation information according to the TCDB (https://www.tcdb.org; Saier et al. 2021) were downloaded using the “Annotations” tab from the MycoCosm website for selected taxonomical ranks, i.e. the phyla Ascomycota and Basidiomycota; subphyla Ustilaginomycotina and Pucciniomycotina; and the order Pucciniales (Supplementary Table S2). The tables were then filtered by selecting the main fungal classes 1 to 5; and auxiliary classes 8 and 9; as well as some specific families and subfamilies within the class 2. In all comparative analyses, the order Pucciniales and subphyla Ustilaginomycotina and Pucciniomycotina were removed from their respective higher taxonomical ranks, referred to thereafter as Other_Pucciniomycotina (i.e. Pucciniomycotina without Pucciniales) and Other_Basidiomycota (i.e. Basidiomycota without Ustilaginomycotina and Pucciniomycotina). The total number of predicted genes was collected for each fungal species. Statistical analyses were conducted with the software R (R Core Team 2018) for comparisons of means between taxonomical groups based on nonparametric analysis (Kruskal-Wallis test); followed by Bonferroni-Dunn’s post hoc test at the threshold level of *P*=0.05.

### Expert annotation of OPT genes

Predicted OPT genes in the order Pucciniales and selected species in the phyla Ascomycota and Basidiomycota were reviewed on the JGI MycoCosm portal using homology and expression information displayed on corresponding tracks. Predicted OPT genes belonging to the different classes of the rust fungi *M. larici-populina* and *Melampsora allii-populina* genomes were manually curated based on support from Sanger- and Illumina-based Expressed Sequence Tags (ESTs) shown on the MycoCosm genome browser, and/or homology to reference OPT genes. Curated OPT were validated by a CD-Search against the Conserved Domain Database (CDD) at the National Center for Biotechnology Information (NCBI, https://www.ncbi.nlm.nih.gov/Structure/cdd/cdd.shtml), and transmembrane domains organization was verified with the TMHMM prediction tool (https://services.healthtech.dtu.dk/service.php?TMHMM-2.0). Additional information regarding genomic organization (e.g. tandem duplicates) was also recorded. Tblastn homology searches with Pucciniales and reference fungal OPT genes were performed against the *Melampsora* spp. genomes to ensure OPT genes were not missed by automatic functional annotations or to verify the absence of OPT-related pseudogenes.

### Phylogenetic analysis of OPTs

OPT protein sequences were collected in representative fungal species from distinct taxonomical groups available in MycoCosm: seven in the order Pucciniales; three in the subphylum Pucciniomycotina (excluding Pucciniales); two in the subphylum Ustilaginomycotina; eight in the subphylum Agaricomycotina (i.e. representing a total of 20 species from the phylum Basidiomycota); and seven species in the phylum Ascomycota. OPT sequences were aligned with MUSCLE (v3.8.1551, Edgar 2004) using default parameters, poorly aligned sequences were trimmed with trimAl (v1.4.1, Capella-Gutierrez et al. 2009) setting the gapthreshold parameter to 0.5. To select the best-fit amino acid substitution models for this trimmed alignment, we used ModelTest-NG (v0.1.6, Darriba et al. 2020). A maximum likelihood inference for our phylogenetic dataset was achieved with RAxML-NG (v0.9.0, Kozlov et al. 2019) using the VT+I+G4 model and 1000 bootstraps replicates. The tree was visualized with FigTree v1.4.4 (http://tree.bio.ed.ac.uk/software/figtree/).

### Transcriptomic resources

Transcriptomic profiling of *M. larici-populina* was established along the life cycle from previously published studies transposed onto the version 2 of the genome (Persoons et al. 2022). Briefly, gene expression data were retrieved for dormant and germinated urediniospores, time course infection of poplar leaves and telia by oligoarrays (Duplessis et al. 2011b; Hacquard et al. 2013) and for basidia, spermogonia and aecia on larch needles by RNAseq (Lorrain et al. 2018a). For this study, we extracted expression profiling of *M. larici-populina* transporter genes from an open transcriptomic dataset (Guerillot et al. 2021; Supplementary Table S3). The Morpheus tool from the Broad Institute website (https://software.broadinstitute.org/morpheus/) was used to present normalized transcript expression levels using the *k*-means partitioning method.

## Results

### Survey of the fungal transportomes reveals a global contraction in the order Pucciniales

Taking advantage of the MycoCosm, we explored the complete predicted transporter catalogs of fungal genomes from the order Pucciniales and other fungi in the Dikarya. Pucciniales exhibit large genomes and gene numbers compared to other fungi; more particularly to other Pucciniomycotina in the same phylum or to Ustilaginomycotina; while other species in the Dikarya present various genome sizes and gene numbers (Aime et al. 2017; Duplessis et al. 2013; Haridas et al. 2020; Miyauchi et al. 2020; Tavares et al. 2014). In order to draw comparisons between different taxonomical ranks, we have chosen to consider the following groups: the order Pucciniales (n=13), Other_Pucciniomycotina (i.e. the subphylum Pucciniomycotina excluding Pucciniales; n=16), the subphylum Ustilaginomycotina (n=28), Other_Basidiomycota (i.e. the phylum Basidiomycota excluding the subphyla Pucciniomycotina and Ustilaginomycotina; n=257), and the phylum Ascomycota (n=737). The total number of genes and transporters for these taxonomical groups are presented in the Figure 1. With an average of 18,103 genes, species of the order Pucciniales presented a significantly larger gene numbers than Other_Pucciniomycotina (mean=7,537), Ustilaginomycotina (mean=6,945) and Ascomycota (mean=11,278), and higher but non-significant gene numbers than in Other_Basidiomycota (mean=15,052) (Supplementary Table S4). Despite these noteworthy differences, the transportome of the Pucciniales exhibited a significant contraction compared to the Ascomycota and Other_Basidiomycota, but not with Other_Pucciniomycotina and Ustilaginomycotina (Figure 1; Supplementary Table S4). The TCDB distributes transporters in seven different classes, the five majors being channels/pores (class 1); electrochemical potential-driven transporters (class 2); primary active transporters (class 3); group translocators (class 4) and transmembrane electron carriers (class 5). According to Saier et al. (2021), classes 8 and 9 correspond to accessory factors involved in transport and incompletely characterized transport systems, respectively. Thus, these classes were collected in Supplementary Table S2 and Supplementary Table S3 for complete information, but were not further considered in our analysis. No significant expansion was detected for any class in the Pucciniales compared to Ascomycota or Other_Basidiomycota (Figure 2; Supplementary Table S4). Significant expansions were only recorded between Pucciniales and Other_Pucciniomycotina (class 3) and Ustilaginomycotina (classes 1, 3 and 5). On the contrary, a significant contraction was observed between Pucciniales and Ascomycota; and between Pucciniales and Ascomycota and Other_Basidiomycota for classes 4 and 2, respectively. Overall, our results indicate an obvious contraction of the transportome in the order Pucciniales compared to Other_Basidiomycota and Ascomycota, despite the larger number of genes observed in their genomes.

**Figure 1.**
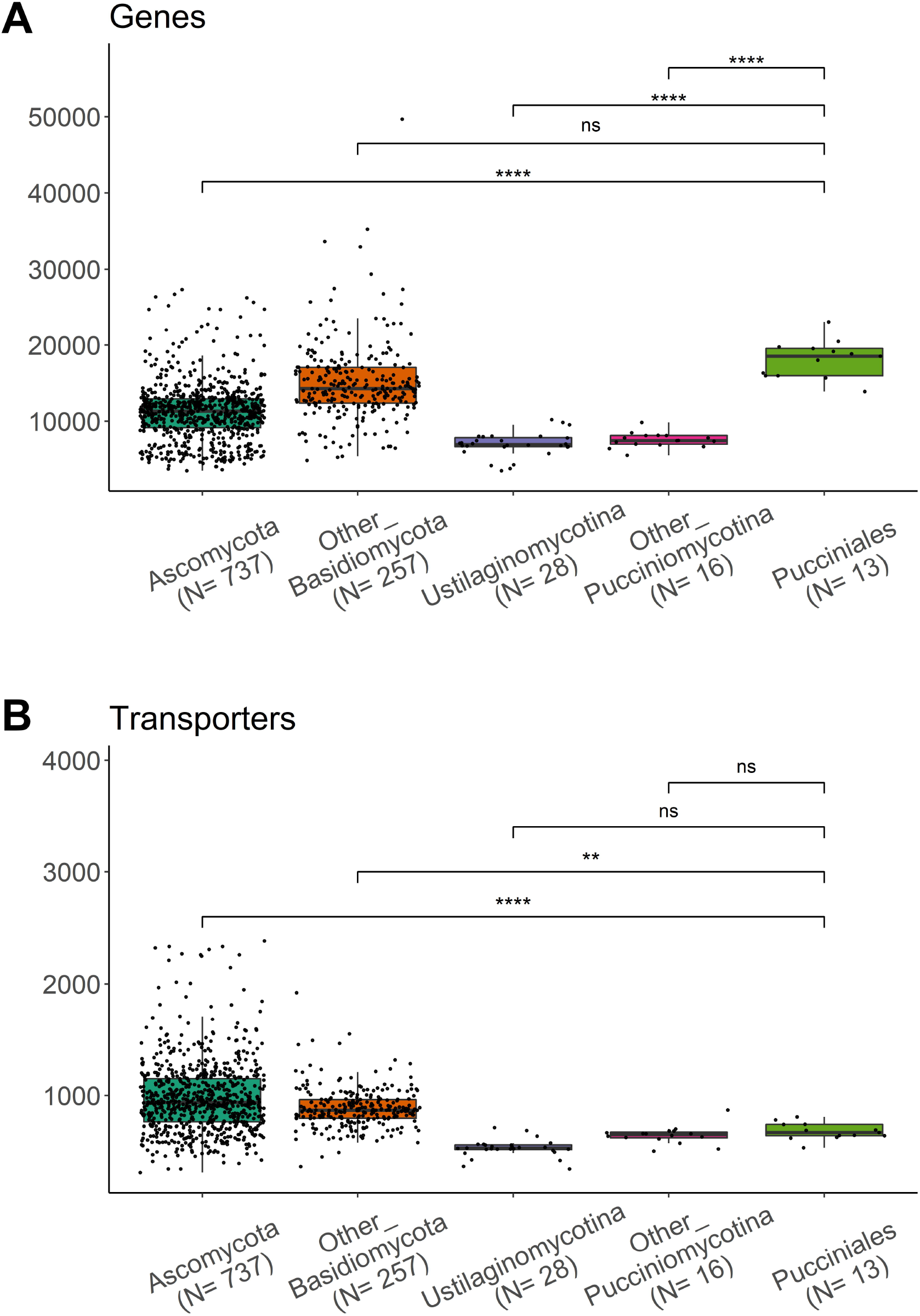
Distribution of number of genes (A) and transporters (B) in fungal genomes. Each dot represents a species. Other_Basidiomycota: Basidiomycota excluding the subphyla Pucciniomycotina and Ustilaginomycotina. Other_Pucciniomycotina: Pucciniomycotina excluding the order Pucciniales. N= number of species considered in each group. Statistical analysis results are presented on top of the box plots (non-parametric Kruskal-Wallis test followed by Bonferroni-Dunn’s post hoc test at the threshold level of *P*=0.05). ns, not significant; **, <0.01; ****, <0.0001. Only comparisons between the order Pucciniales and the other taxonomical groups are presented.

**Figure 2.**
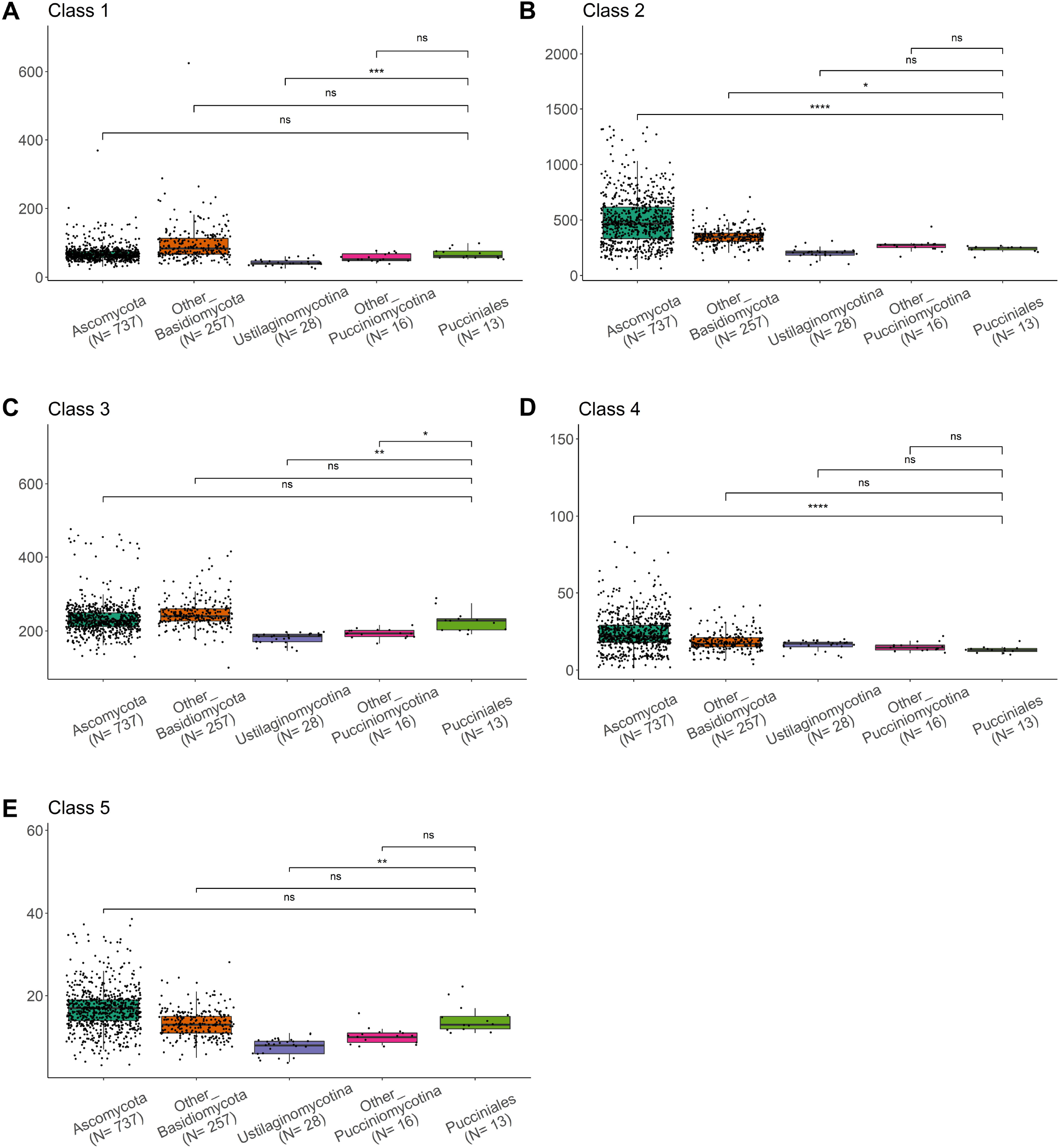
Distribution of number of transporter genes in the five main TCDB classes. A, class 1 (channels/pores); B, class 2 (electrochemical potential-driven transporters); C, class 3 (primary active transporters); D, class 4 (group translocators) and E, class 5 (transmembrane electron carriers). Each dot represents a species. Other_Basidiomycota: Basidiomycota excluding the subphyla Pucciniomycotina and Ustilaginomycotina. Other_Pucciniomycotina: Pucciniomycotina excluding the order Pucciniales. N= number of species considered in each group. Statistical analysis results are presented on top of the box plots (non-parametric Kruskal-Wallis test followed by Bonferroni-Dunn’s post hoc test at the threshold level of *P*=0.05). ns, not significant; *, <0.05; **, <0.01; ***, <0.001; ****, <0.0001. Only comparisons between the order Pucciniales and the other taxonomical groups are presented. TCDB, transporter classification database.

### Survey of class 2 transporter families reveals specific expansions of OPT and metal transporters

We scrutinized in particular the electrochemical potential-driven transporters (TCDB class 2), which contains transporters previously described in rust fungi. Among Dikarya, the porters (TCDB subclass 2.A Porters: uniporters, symporters, antiporters) represented more than 99% of the transporters from the class 2 (Supplementary Table S2). Among them, the major facilitator family (MFS; TCDB family 2.A.1) which contains HXT1 reported in *U. fabae* and *P. striiformis* f. sp. *tritici* (TCDB subfamily 2.A.1.1) showed a significant contraction in the Pucciniales compared to Ascomycota and Other_Basidiomycota (Figure 3A; Supplementary Table S4). The amino acid-polyamine-organocation family (APC; TCDB 2.A.3 family), which corresponds to the AATs from *U. fabae* presented a significant contraction in the Pucciniales compared to Ascomycota (Figure 3B; Supplementary Table S4). Considering the overall contraction observed in the class 2 for Pucciniales, we searched for potential expansions at the family level. A total of 12 families showed a significant expansion in the Pucciniales compared to Ascomycota and/or Other_Basidiomycota (Supplementary Table S4) and only four families presented particularly remarkable expansions (Figure 3C-F). The most dramatic expansion was observed in the OPT (TCDB family 2.A.67) with an average of 25 OPTs in the Pucciniales and 9, 14, 7, 7 OPTs in the Ascomycota, Other_Basidiomycota, Ustilaginomycotina and Other_Pucciniomycotina, respectively (Figure 3E). The three other families were all related to metal transport: the cation diffusion facilitator family (CDF; TCDB family 2.A.4) ensuring heavy metal transport, the metal ion (Mn2+ -iron) transporter family (Nramp; TCDB family 2.A.55), and the vacuolar iron transporter (VIT; TCDB family 2.A.89) (Figure 3). Other families presented significant expansions with Other_Pucciniomycotina and/or Ustilaginomycotina but not with other fungi in the Dikarya (Supplementary Table S4). Our results show that despite a global reduction of the Porters class, dramatic expansions can be retrieved in the Pucciniales lineage for transport systems related to metals and oligopeptides, highlighting putative innovations in the evolution of rust fungi.

**Figure 3.**
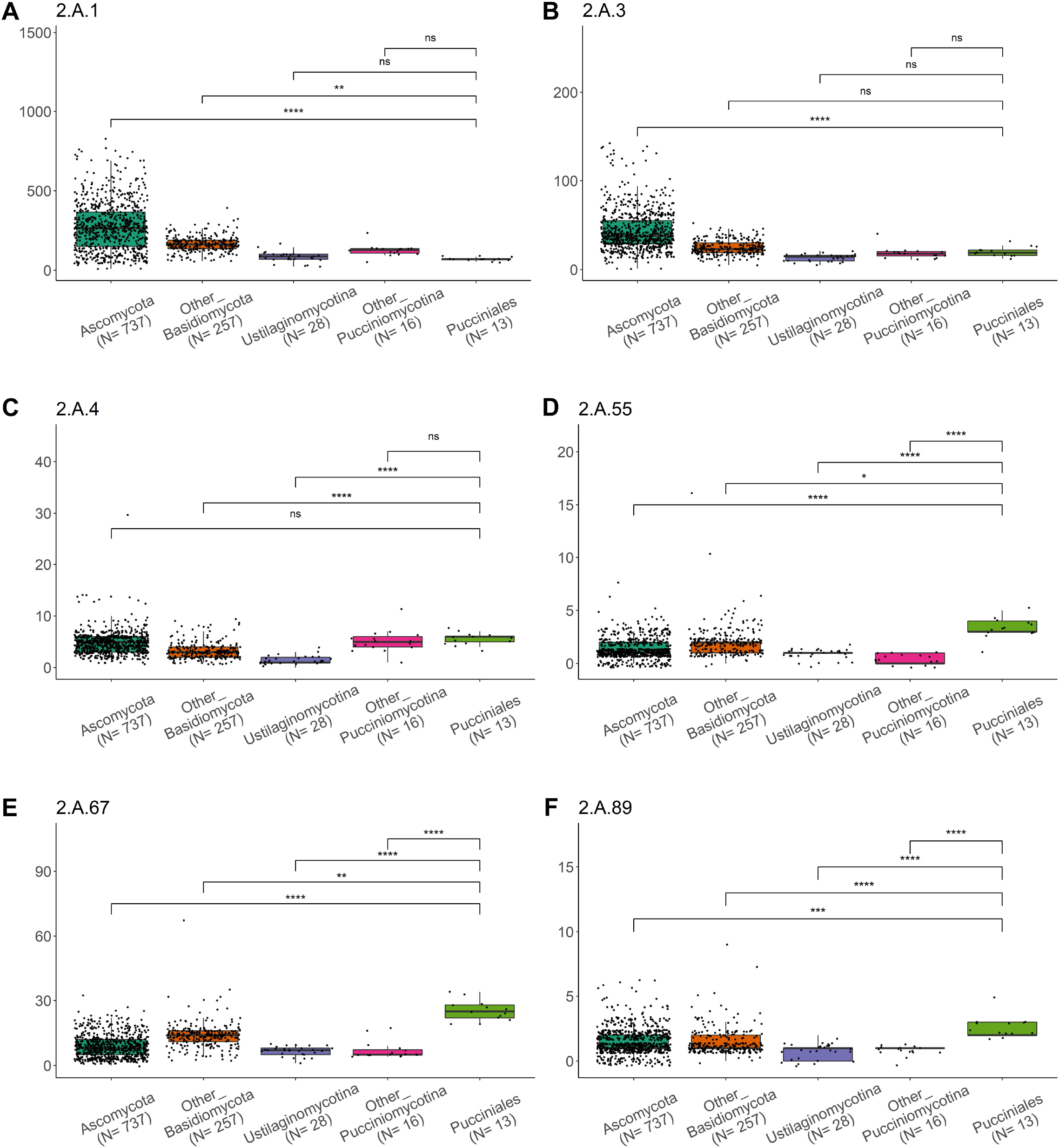
Distribution of number of transporter genes in six selected TCDB families. A, 2.A.1 (the major facilitator superfamily, MFS); B, 2.A.3 (the amino acid-polyamine-organocation family, APC); C, 2.A.4 (the cation diffusion facilitator family, CDF); D, 2.A.55 (the metal ion transporter family, Nramp); E, 2.A.67 (the oligopeptide transporter family, OPT) and F, 2.A.89 (the vacuolar iron transporter family, VIT). Each dot represents a species. Other_Basidiomycota: Basidiomycota excluding the subphyla Pucciniomycotina and Ustilaginomycotina. Other_Pucciniomycotina: Pucciniomycotina excluding the order Pucciniales. N= number of species considered in each group. Statistical analysis results are presented on top of the box plots (non-parametric Kruskal-Wallis test followed by Bonferroni-Dunn’s post hoc test at the threshold level of *P*=0.05). ns, not significant; *, <0.05; **, <0.01; ***, <0.001; ****, <0.0001. Only comparisons between the order Pucciniales and the other taxonomical groups are presented. TCDB, transporter classification database.

### Annotation survey of OPTs confirms their expansion in the order Pucciniales

In order to better define the evolution of rust OPTs, we first performed a manual curation of OPT genes in the poplar rust fungus *M. larici-populina*, a model rust species for which extensive transcript resources are available for expert annotation. A total of 23 OPT genes were predicted in the genome of *M. larici-populina*, of which 17 needed curation (Table 1). With one exception (fusion of two alternative gene models), an alternative best gene model could systematically be retrieved from the JGI MycoCosm at the loci of OPT gene which needed curation (Table 1). OPT genes exhibited a complex structure with an average of 26 exons (9-36 exons; Table 1) and most of the curations were exon additions (12) or exon corrections (4). Almost all curations were based on the presence of expression support (RNAseq coverage track and Sanger or RNAseq EST tracks from MycoCosm genome browser). One model needed a fusion between alternate models due to a long intron of 2.5Kb generated by a transposable element insertion. Only three gene models were clustered within < 50 Kb on the linkage group LG_12 (Table 1). tblastn homology searches did not identify other OPT genes or pseudogenes in the genome. A similar expert curation approach was carried out in *M. allii-populina*, a close relative species also causing poplar rust disease. A total of 28 genes were predicted, of which 15 needed curations with exon additions or curations (Supplementary Table S5). Only four genes were found in clusters, with two tandem duplications lying on the scaffold_88. OPT genes from other published Pucciniales genomes were also considered, but not manually curated due to the lack of extensive expression support. Nevertheless, the number of exons and predicted TMs were collected in order to identify genes which obviously need attention (Supplementary Table S6). The manual curation performed in the two species from the family Melampsoraceae confirms the OPT genes expansion in rust fungi and the absence of extensive tandem duplications and gene clusters suggests that these genes emerged from old duplication events.

**Table 1.**
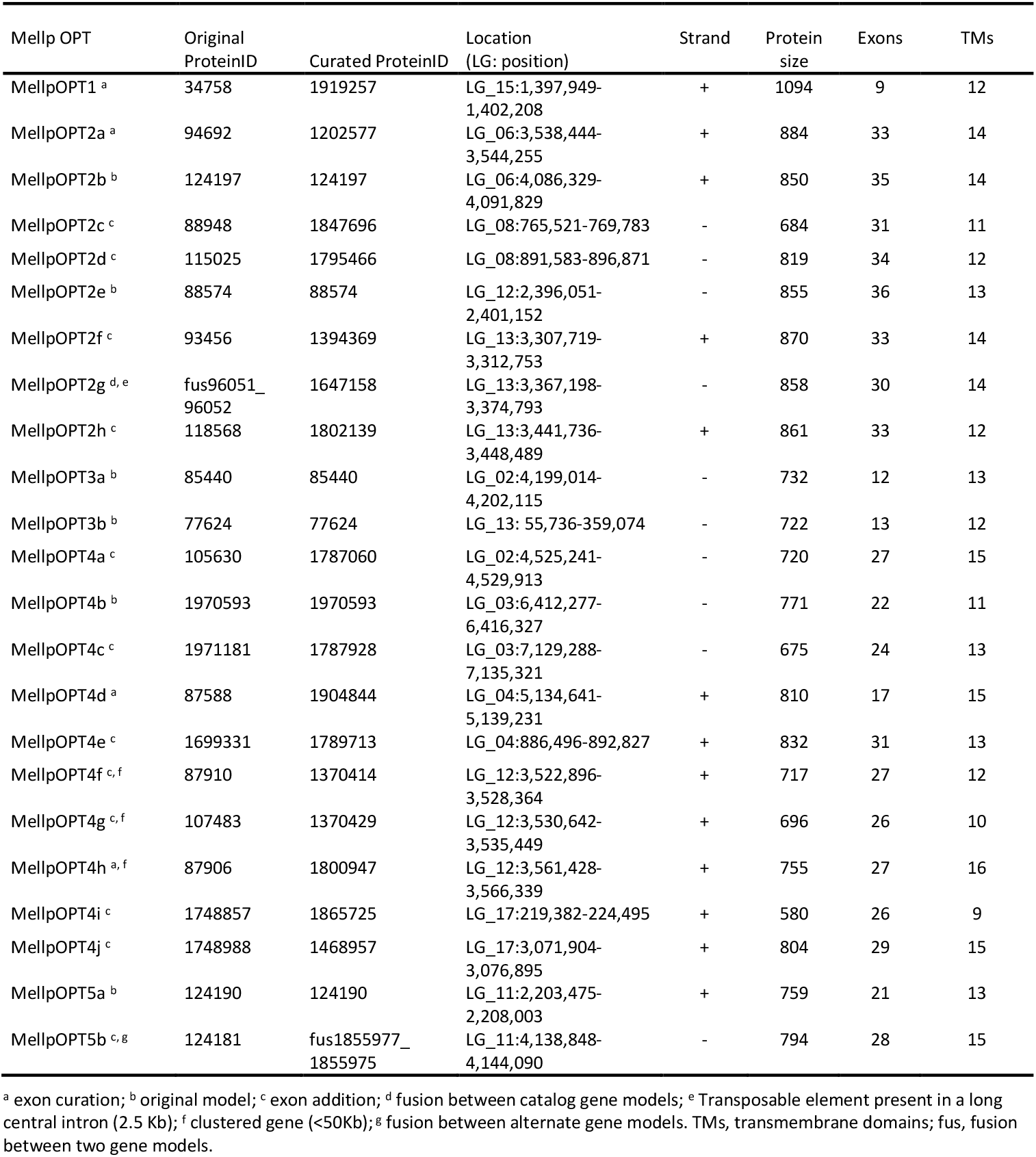
Annotation of the *Melampsora larici-populina* oligopeptide transporter (OPT) gene family.

### Phylogenetic analysis of OPTs reveals several expansions at the root of the order Pucciniales

We conducted a phylogenetic analysis in order to obtain a better knowledge of the evolution of the OPT family in rust fungi. We collected 396 OPTs from a selection of 27 representative fungal species in the Dikarya (Table 2). Species were selected in the different taxonomical groups with the *a priori* criterion of interaction with plants in their trophic mode and the selected fungi were either pathogens, mutualistic symbionts or wood-decayers, except for *Sporobolomyces roseus* in the Other_Pucciniomycotina, a unicellular “red” yeast which was added for its taxonomical proximity to Pucciniales. OPTs previously described in three additional Ascomycota species were added as references (Table 2). The selected species were not outliers in their taxonomical categories regarding their gene numbers, with the two exceptions of *Microbotryum lychnidis-dioicae* which displayed 16 OPTs (mean=7 in Other_Pucciniomycotina) and *Lactarius quietus* which displays 67 OPTs (mean=14 in Other_Basidiomycota). Of the 396 OPTs, 361 sequences were retained after removing truncated OPTs and processing trimming on the alignment. The Figure 4 shows the unrooted phylogenetic tree generated with RAxML for the 361 remaining fungal OPTs (Supplementary Table S7). A total of 19 reference OPTs previously functionally studied in the ascomycetes *Aspergillus fumigatus, Candida albicans* and *Saccharomyces cerevisiae* were used to define clades in the tree. Major nodes holding branches with a bootstrap value > 50% were also considered to define clades. Fungal OPTs were distributed among six major clades. The different taxonomical groups were distributed in all OPT clades except for clades 5 and 6, which contains species from the Pucciniomycotina and the Ustilaginomycotina only; and species from the Ascomycota only, respectively (Figure 4). The clade 1 showed a dichotomy between Ascomycota and Basidiomycota species, particularly, one major branch holding a *L. quietus* specific expansion also carried OPTs from the Other_Basidiomycota group only. The major expansion seen in the outlier species *L. quietus* likely emerged from recent duplications since 75 % of the sequences grouped along the same branch and were encoded by several genes organized in tandem in the genome (Figure 4; Supplementary Table S6). OPTs from the Pucciniales were found in all clades except from the clade 6. Also, three expansions specific to the Pucciniales were present in clades 2, 4 and 5. Noteworthy, the most remarkable expansion was present on a major branch of the clade 4 and accounted for 53% of the Pucciniales OPTs. In each of the Pucciniales expansion, OPTs were distributed in different branches with representatives from the different taxonomical families Coleosporiaceae, Melampsoraceae and Pucciniaceae, suggesting ancient duplication events (Supplementary Figure S1). The plant pathogen *M. lychnidis-dioicae* from the Pucciniomycotina also presented an expansion in the clade 4, however it was found on a branch disconnected from the major specific expansion noted for the Pucciniales, indicating these OPTs likely arose from independent old duplication events (Figure 4). To conclude, this phylogenetic analysis indicates that the OPT expansion likely occurred through ancient duplications at the root of the order Pucciniales, then subsequently in specific taxonomical families in the course of the evolution. This result supports both a diversification of the OPT gene family and possibly some level of specialization in specific OPT clades.

**Figure 4.**
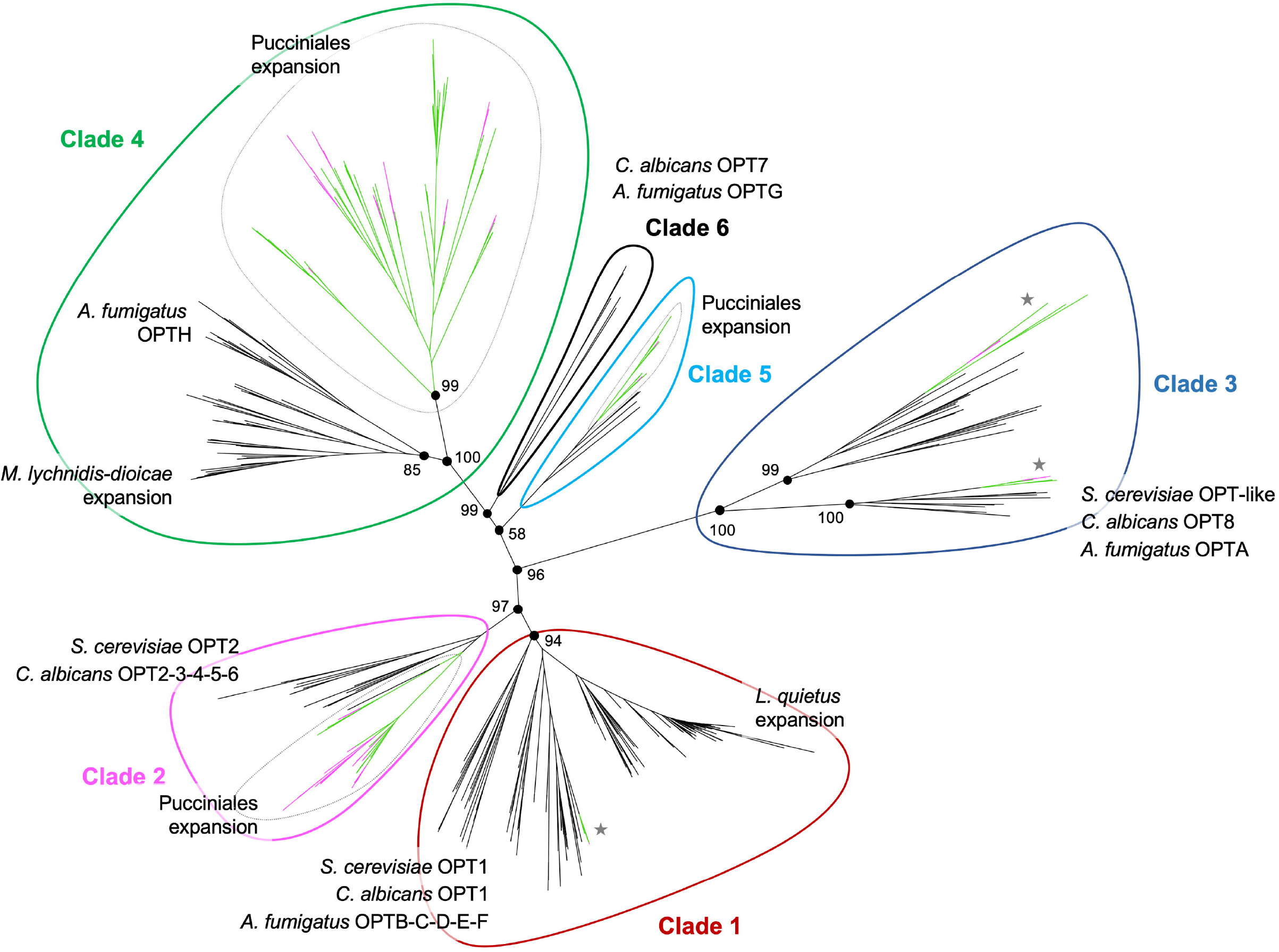
Phylogenetic relationships among the oligopeptide transporter (OPT) gene family in selected ascomycetes and basidiomycetes. A tree was built based on 361 OPT protein sequences using the maximumlikelihood method with RAxML-NG program. Bootstrap values above 50 (percentage of 1000 replicates) and supporting a node used to define clusters are indicated. Clades were arbitrarily defined from 1 to 6 based on the presence of reference OPTs previously described in *Aspergilus fumigatus, Candida albicans* and *Saccharomyces cerevisiae*. Branch lengths are proportional to genetic distances. Reference OPTs are indicated at proximity of the clade which contains them. Remarkable expansions observed in the order Pucciniales and in the species *Microbotryum lychnidisdioiceae* and *Lactarius quietus* are mentioned. Pucciniales OPT expansions in clades 2, 4 and 5 are marked by dotted lines. Branches with Pucciniales OPT in clades 1 and 3 are marked by a star.

**Table 2.** Number of oligopeptide transporter (OPT) genes in selected fungal species used in the phylogenetic analysis.

| Species | ID<sup>a</sup> | Taxonomy | OPT number <sup>b</sup> |
| --- | --- | --- | --- |
| *Aspergillus fumigatus* <sup>d</sup> | Afu | Ascomycota | 8 (8) |
| *Blumeria graminis* f. sp. *hordei* <sup>c</sup> | Blugr | Ascomycota | 5 (5) |
| *Candida albicans* <sup>d</sup> | Canalb | Ascomycota | 8 (8) |
| *Leptosphaeria maculans* <sup>c</sup> | Lepmu | Ascomycota | 3 (3) |
| *Magnaporthe oryzae* <sup>c</sup> | Magor | Ascomycota | 12 (12) |
| *Saccharomyces cerevisiae* <sup>d</sup> | Sacce | Ascomycota | 3 (3) |
| *Zymoseptoria tritici* <sup>c, i</sup> | Zymtr | Ascomycota | 8 (8) |
| *Armillaria mellea* <sup>c</sup> | Armme | Other_Basidiomycota | 18 (16) |
| *Heterobasidion annosum* <sup>c</sup> | Hetan | Other_Basidiomycota | 11 (11) |
| *Laccaria bicolor* <sup>e</sup> | Lacbi | Other_Basidiomycota | 9 (9) |
| *Lactarius quietus* <sup>e</sup> | Lacqui | Other_Basidiomycota | 67 (50) |
| *Phanerochaete chrysosporium* <sup>f</sup> | Phchr | Other_Basidiomycota | 17 (17) |
| *Piriformospora indica* <sup>g</sup> | Pirin | Other_Basidiomycota | 7 (7) |
| *Pisolithus microcarpus* <sup>e</sup> | Pismi | Other_Basidiomycota | 11 (9) |
| *Russula emetica* Přilba <sup>e</sup> | Ruseme | Other_Basidiomycota | 10 (9) |
| *Microbotryum lychnidis-dioicae* <sup>c</sup> | Micld | Other_Pucciniomycotina | 16 (13) |
| *Mixia osmundae* <sup>c</sup> | Mixos | Other_Pucciniomycotina | 4 (4) |
| *Sporobolomyces roseus* <sup>h</sup> | Sporo | Other_Pucciniomycotina | 5 (5) |
| *Cronartium quercuum* f. sp. *fusiforme* <sup>c</sup> | Croqu | Pucciniales | 18 (17) |
| *Melampsora allii-populina* <sup>c</sup> | Melap | Pucciniales | 28 (26) |
| *Melampsora larici-populina* <sup>c</sup> | Mellp | Pucciniales | 23 (23) |
| *Melampsora lini* <sup>c</sup> | Melli | Pucciniales | 19 (18) |
| *Puccinia graminis* f. sp. *tritici* <sup>c</sup> | Pucgr | Pucciniales | 26 (22) |
| *Puccinia striiformis* f. sp. *tritici* <sup>c</sup> | Pucst | Pucciniales | 21 (20) |
| *Puccinia triticina* <sup>c</sup> | Puctr | Pucciniales | 23 (22) |
| *Sporisorium reilianum* <sup>c</sup> | Spore | Ustilaginomycotina | 8 (8) |
| *Ustilago maydis* <sup>c</sup> | Ustma | Ustilaginomycotina | 8 (8) |

<sup>a</sup> ID, MycoCosm species abbreviations (used in the phylogenetic dataset); <sup>b</sup> Number of genes encoding OPT in the corresponding fungal species (number of OPT sequences retained in the phylogenetic analysis); <sup>c</sup> pathogen; <sup>d</sup> fungal species used as OPT reference; <sup>e</sup> mutualistic symbiont; <sup>f</sup> wood decayer; <sup>g</sup> versatile mutualistic symbiont; <sup>h</sup> unicellular red yeast selected for taxonomical proximity; <sup>i</sup> heterotypic synonym, *Mycosphaerella tritici*.

### Transcript profiling along the life cycle of the poplar rust fungus *M. larici-populina* highlights specific expression profiles of transporter genes

*M. larici-populina* is a model rust fungus with extensive transcriptome information acquired along its whole lifecycle allowing for the study of expression patterns in given cellular categories (Guerillot et al. 2021; Louet et al. 2021). We extracted normalized expression data for 684 transporter genes, out of 743 total transporters for this rust fungus (missing data are from early oligoarraybased transcriptome studies; Supplementary Table S3). The Figure 5 presents a condensed view of the general expression profile of 547 transporter genes falling in the TCDB classes 1 to 5 out of 592 (92%) sorted in 10 k-means (expression data for classes 8 and 9 was also collected and is shown in Supplementary Table S3). An expanded version of this expression profile for the 684 transporter genes, including the classes 8 and 9, is presented sorted by transporter subfamilies or sorted by their k-means in the Supplementary Figure S2A and S2B, respectively. Overall, only a limited number of transporters showed expression mostly restricted to a single stage, such as in the k-means k2 (poplar in plant biotrophic growth), k4 (larch) and k6 (basidia) (Figure 5). Many transporter genes presented a high level of expression in urediniospores (spores infecting the poplar host) associated with moderate or high levels of expression in the host plant poplar in k-means k10 (19 %) and k7 (17 %), respectively; or with expression in basidia (producing spores infecting the alternate host, larch) in k-mean k5 (11 %). Finally, several transporter genes showed peaks of expression at early, intermediate or late stages of poplar leaf infection in k-means k3; k2 and k9; and k1, respectively (Figure 5). These results indicate a highly dynamic regulation of transporter genes expression along plant infection and some level of expression specificity in one host or another. When scrutinized into details, no complete transporter family exhibited a specific expression for all of its gene members at a single stage, although a few gene families may show a preferential expression at a given stage for a majority of their gene members (Supplementary Figure S2A). Metal-related gene families expanded in rust genomes showed various patterns of expression in *M. larici-populina* spores (either in basidia or in urediniospores) or during host infection, with a preferential expression in the later situation (Supplementary Figure S2A). The survey of the OPT gene family showed that all genes are expressed at least at a given stage, with nine genes showing a high expression in urediniospores and four showing a peak of expression in basidia (Supplementary Figure S2A). Interestingly, while three and nine OPT genes showed a specific peak of expression at 24 and 96 hours postinoculation in infected poplar leaves, only a single OPT gene showed a peak of expression in spermogonia and aecia formed on the alternate host, larch. However, a few other OPT genes showed lower levels of expression on both host plants. In total, 10 out of 24 OPT genes were specifically expressed *in planta* suggesting a specific transport role during the biotrophic growth. To conclude, profiling of *M. larici-populina* transporter genes shows dynamic patterns of expression regulation in spores infecting both host plants with some specificity within transporter families toward one host or the other for some gene members, but not over a complete transporter family. Particularly, the OPT genes are expressed at all stages surveyed in the poplar rust fungus and almost half of them show a specific expression during host infection, either in poplar or in larch.

**Figure 5.**
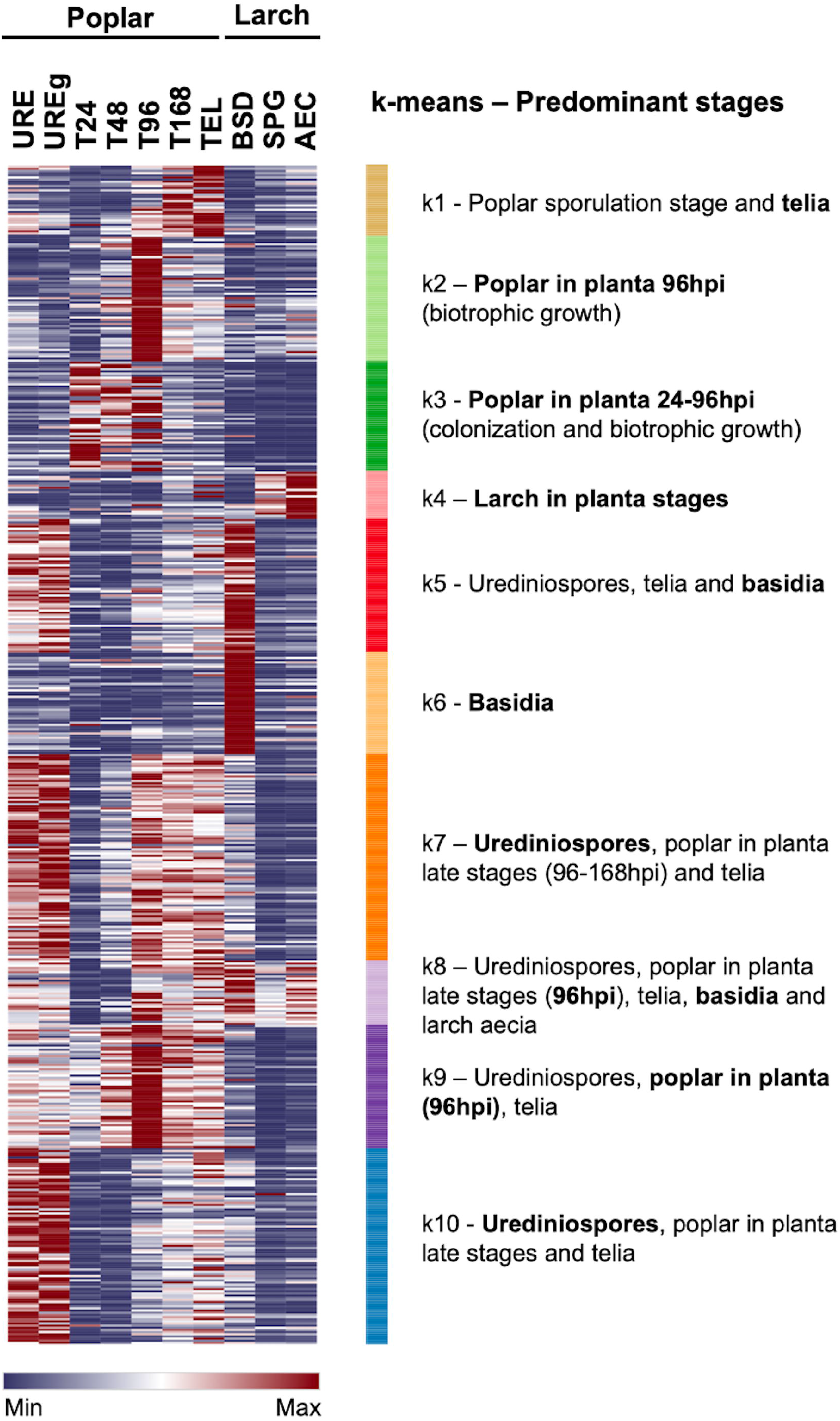
Expression profiling of the poplar rust fungus transporter gene repertoire across its complete life cycle. Expression levels of 547 transporter genes were collected from various life stages and sorted by k-means partitioning (k= 10). Life stages are presented on top according to the poplar rust life cycle (infection of the telial host poplar followed by infection of the aecial host larch). Color scale from blue to red represents normalized expression levels from 0 to maximum value at each stage. Biological stages showing high expression levels (i.e., above average values) in each k-mean are depicted on the right side of the profile and the predominant stage, at which expression peaks, is highlighted in bold. URE and UREg, dormant and germinating poplar rust urediniospores, respectively; T24 to T168, hours post-inoculation in a time course infection series on poplar leaves inoculated with urediniospores; TEL, telia; BSD, basidia; SPG and AEC, spermogonia and aecia formed on larch needles, respectively.

## Discussion

Rust fungi are obligate biotrophic fungal plant pathogens that strictly depend on their hosts to feed, grow and reproduce (Voegele and Mendgen 2011). Early molecular studies of rust infection structures such as the haustorium pointed out the specific expression and localization of a suite of transporters contributing to fungal nutrition during the biotrophic growth (Lorrain et al. 2019; Voegele et al. 2009). However, our knowledge about transport systems in rust fungi remains limited. At the time of the publication of the first rust fungal genomes *M. larici-populina* and *P. graminis* f. sp. *tritici*, a survey of transporter genes was carried out with limited possibilities in term of comparison with only a handful of fungal genomes available (Duplessis et al. 2011a). Here, we performed an annotation survey of the transportome of rust fungi and a comparison within the Dikarya by harvesting genomic data from more than a thousand fungi. We combined this genomic analysis with transcriptomic profiles in the model rust fungus *M. larici-populina* along its entire life cycle. Our analysis reveals striking evolutionary trends in the transport systems of rust fungi with several gene family expansions. These genes show dynamic expression patterns during host infection and other stages of the rust biological cycle. Altogether, our results highlight some specificities likely related to the biotrophic status of rust fungi and opens interesting perspectives to better comprehend nutrients acquisition in these plant pathogens.

Overall, the transportome of fungi in the Dikarya represents on average 7.8% of the total gene content. This proportion greatly varies between higher taxonomical ranks due to the genome size and the number of genes predicted in these genomes. With their large gene numbers, genomes of the order Pucciniales are marked by a dramatic reduction of their repertoire of transporter genes compared to other Dikarya. This tendency is also observed in the entire subphylum Pucciniomycotina, to which Pucciniales belong, and in the subphylum Ustilaginomycotina, which exhibits at the contrary compact genomes with a reduced number of genes (Duplessis et al. 2013; Kamper et al. 2006; Ullmann et al. 2022). Such a global contraction seen in Pucciniales and in some Basidiomycota subphyla likely reflects a specialization in their trophic modes, e.g., towards biotrophy, or adaptation to specific ecological niches (Aime et al. 2014; Spatafora et al. 2017). This transportome reduction in rust fungi is mostly due to contractions of transporter families falling in the class 2 (electrochemical potential-driven transporters) and class 4 (translocators), which altogether accounts for 45% of all rust transporters. Rust fungi fully depend on their host plant for the acquisition of carbon and nitrogen, and hexose or amino acids were shown to be preferentially imported from haustoria during their biotrophic growth (Struck 2015; Voegele et al. 2009). Isolation of RNA from haustorial infection structures of the bean rust fungus *U. fabae* identified genes highly expressed in these specific biotrophic features (Hahn and Mendgen, 1997). Among those genes, a suite of transporters -i.e. the hexose transporters Uf-HXT1 of the 2.A.1.1 subfamily and Uf-AAT1, Uf-AAT2 and Uf-AAT3 of the 2.A.3 family-showed specific and high expression patterns and/or could be localized at the periphery of these feeding haustorial structures (Hahn et al. 1997; Struck et al. 2002, 2004; Voegele et al. 2001; Voegele and Mendgen 2011). The high expression of orthologs of these transporter genes was confirmed in other rust fungi by transcriptomic approaches, indicative of a conserved transport set up in these biotrophic pathogens (Dobon et al. 2016; Duplessis et al. 2011a, 2011b; Garnica et al. 2013). Interestingly, the two families 2.A.1 and 2.A.3 are under contraction in rust fungi and only a few gene members are present in each of these families. A survey of their expression along the life cycle of the poplar rust fungus confirms their high expression in both alternating host plants with some specificities on one host or the other. Their high expression, as well as their contraction, highlight similar evolutionary trends towards specialization during biotrophic growth. A high expression was similarly reported in the mutualistic biotrophic fungus *Laccaria bicolor* during the interaction with a host plant, however some amino acid transporters subfamilies are marked by moderate expansions, highlighting different adaptation strategies among mutualistic and pathogenic biotrophs (Lucic et al. 2008).

Metals such as iron or divalent metal ions are essential for fungal growth. However, they may also represent a source of toxicity under accumulation, and they require a fine tuning of their homeostasis (Robinson et al. 2021). Specific transporter families are adapted for the import, export or storage of such metals and contribute to metal homeostasis. Metal homeostasis and regulation of metal transport also play a role in virulence and during the interaction between fungi and host plants (Gerwien et al. 2018; González-Guerrero et al. 2016). The systematic comparison of transporter families in Pucciniales against other Dikarya revealed significant expansions in rust fungi for three gene families which are related to metal transport. The family 2.A.4 (CDF) allows export of divalent cations from the cytoplasm, whereas the family 2.A.55 (Nramp) allows the import of iron and the family 2.A.89 (VIT) allows the sequestration of iron in the vacuole (González-Guerrero et al. 2016; Sorribes-Dauden et al. 2020). Beyond expansion, these gene families also show dynamic patterns of expression across the life cycle of the poplar rust fungus. Particularly, the Nramp transporters involved in iron import are showing a preferential expression during infection of the poplar host. These observations suggest that rust fungi might scavenge metals such as iron from their host during the colonization process and it would be interesting to determine more precisely whether the import takes place in the haustorial structures, i.e. from the invaded host cell cavities; or in infection hyphae progressing in the apoplastic space. It would be also interesting to determine whether the CDF are related to metal tolerance while progressing in the host leaf tissue or to metal homeostasis through maintenance of a balance between import and export of metals for instance. Although the role of metal transport in fungi has been studied during symbiosis or virulence (Gerwien et al. 2018; González-Guerrero et al. 2016), nothing is known in rust fungi, which opens interesting perspectives in understanding the role of metal homeostasis in rust fungal biology.

Genomic studies have revealed that rust fungi lack key genes for direct assimilation of inorganic nitrate and sulfur, which indicates that they need to acquire these nutrients under other forms through specific transport systems (Duplessis et al. 2011a). As pointed out above, amino acids were shown to represent a major source of nitrogen acquired from the host through adapted transporters in haustoria (Struck 2015). A large number of OPT genes was noticed in early rust genomic projects and high expression of OPT genes was also observed in isolated haustoria of the wheat stem rust fungus, suggesting that these transporters may represent an additional way to derive nitrogen and also sulfur from the infected host (Duplessis et al. 2011a; Garnica et al. 2013). OPTs are transport systems localized at the plasma membranes and involved in the import of oligopeptides (Lubkowitz 2011). OPTs have been particularly studied in a few model ascomycetes in which gene numbers varied from three to eight genes; and the encoded OPTs showed transport preferences for different oligopeptide forms (Becerra-Rodriguez et al. 2020; Hartmann et al. 2011; Hauser et al. 2000; Reuß and Morschhäuser 2006). Although testing all oligopeptide combinations cannot be reasonably achieved (Lubkowitz 2006), some OPTs showed a high affinity for glutathione which is a modified tripeptide, whereas others have preferential affinity for tetra- or penta-peptides, and other OPTs are able to transport oligopeptides of up to eight residues (Becerra-Rodriguez et al. 2020). In *S. cerevisae* wine strains which possess a specific set of fungal OPTs (FOTs) acquired through horizontal gene transfer, substrate specificities could also be demonstrated in mutants expressing single FOT genes, suggesting that sequence divergence can underlie di- and tri-peptide preferences (Becerra-Rodriguez et al. 2021). Our comparative study identified several OPT genes expansions in different species of the phylum Basidiomycota, with specific and prominent expansions at the root of the order Pucciniales. Moderate expansion of OPT genes has been previously reported in the ectomycorrhizal fungus *L. bicolor*, and some members are particularly expressed during the biotrophic interaction with the host plant (Lucic et al. 2008). Furthermore, the obligate biotrophic pathogen *M. lychnidis-dioicae* in the Pucciniomycotina presents an expansion of OPT genes in its genome (Perlin et al. 2015), some of which may be related to the adaptation to the host plant (Badouin et al. 2017). *M. lychnidis-dioicae* OPT genes have been shown to be particularly expressed during the late interaction with the host plant, suggesting a possible role in the fungus nutrient acquisition and interestingly, also in nutrient-limited conditions (Perlin et al. 2015; Toh et al. 2017). Here, the OPT genes expansion is the most remarkable in rust fungi despite the overall transportome reduction. On the one hand, several OPT genes are conserved among species of the Dikarya and may share substrate specificities and eventually common regulatory elements related to the sensing and transport of amino acids or nitrogen. On the other hand, the large expansions shared by Pucciniales in OPT clades 2 and 4 might underlie specific mechanisms for acquiring nitrogen and sulfur in the limiting condition of biotrophic growth inside the host plant through these transport systems. Limited tandem duplications of OPT genes were noticeable in the genomes of Pucciniales suggesting that these duplications are rather ancient and might have arisen early on during the radiation of the Pucciniales and before the separation of taxonomical families (Aime et al. 2018). Additional expansions have emerged in the OPT clade 4 more recently in the families Melampsoraceae and Pucciniaceae, indicating that gene expansion in this transporter family occurred through successive duplication events. Interestingly, apart from these OPT expansions in the order Pucciniales, only a limited number of striking expansions were noticed in other species. The expansion of OPT genes in the genomes of rust fungi and of anther smuts in the subphylum Pucciniomycotina has recently been suggested as a possible convergent adaptation to the parasitic lifestyle (Zhou et al. 2022). Apart from the expansion reported in *M. lychnidis-dioicae*, the basidiomycete *L. quietus*(Agarycomycotina; Russulales) presents the most dramatic expansion in fungi with more than 60 genes which emerged mostly through rounds of recent tandem duplications in the OPT clade 1 (Miyauchi et al. 2020). No other close-relative of *L. quietus* in the Russulales showed any particular expansion of OPT genes. *L. quietus* is an ectomycorrhizal symbiont of oak trees with a high level of host specificity, which acquire nutrients in forest soil notably through the release of secreted enzymes (Courty et al. 2007). It would be particularly interesting to test whether OPT genes from this biotrophic mutualistic fungus are all active and expressed, and whether they may be related either to the interaction with the host and/or nutrient gathering in the form of oligopeptides in the surrounding soil.

Rust OPT genes were deemed highly expressed in planta in different rust transcriptomic studies (Duplessis et al. 2011a; Garnica et al. 2013; Lorrain et al. 2019). Interestingly, OPT genes do not show constitutive expression along the whole life cycle of the poplar rust fungus, at the contrary, they are either expressed in the spores infecting each alternate host, or directly during biotrophic growth inside the hosts. The high expression of a series of OPT in isolated haustoria of the wheat stem rust, along with the expression of amino acid transporters clearly confirm that this specialized structure plays a key role as a nutritional center in the course of interaction with the host plant (Garnica et al. 2013; Lorrain et al. 2019). Transcriptomic profiling of host-infection in the maize pathogen *U. maydis* pointed out that three OPT genes were expressed in a specific infection-related cluster associated to biotrophy (Lanver et al. 2018). The systematic mutation of these three genes - i.e., single to triple mutants generated by Crispr/Cas9 - impaired fungal virulence and complementation with any of the genes restored virulence. Previous studies have demonstrated that *U. maydis* mutants can be successfully complemented by Pucciniales genes (Hu et al. 2007). It would be interesting to perform such a complementation to determine whether the closest rust OPT homologs could restore virulence and also to test if other specific Pucciniales OPT could do so. Our results show that only a limited fraction of the OPT genes are expressed in the alternate host of the poplar rust fungus. It would be particularly interesting to determine the extent of the OPT genes which are expressed in haustoria formed in the alternate host plant, and whether there are specific OPT genes adapted to a given host. A major question is: what is the source of oligopeptides transported by the rust fungi during the interaction with their hosts? The rust haustorium is surrounded by an extra-haustorial matrix which is composed of both fungal and host material (Voegele and Mendgen 2003). Numerous genes encoding proteases have been shown to be expressed during host colonization (Duplessis et al. 2011b). Various extracellular proteases -including metalloproteases-were also identified in infection structures of the bean rust *U. fabae* analyzed *in vitro* (Rausher et al. 1995). These proteases may release oligopeptides from degrading proteins residing in this compartment. It would be particularly interesting to attempt at determining the flux of oligopeptides which is derived from the host plant and what is the material acting as a source for such oligopeptides. Although the most probable role of OPT remains the transport of oligopeptides, it cannot be excluded that some specific OPT may relay the perception of signals in the course of the interaction. In plants, some OPTs are involved in the transport of metal-chelates (Lubkowitz 2006). Considering the expansion of metal transporter families in rust fungi, it would be interesting to determine if some Pucciniales-specific OPTs could play a similar role in rust fungi.

In conclusion, we have shown that the transportome of fungi in the order Pucciniales is marked by gene family expansions related to metal transport and most remarkably to oligopeptide transport. Members of these expanded families are expressed at specific stages of the rust life cycle and more particularly during host infection. OPTs represent a new route for acquiring nitrogen and sulfur from the host during the biotrophic growth. Thus far, amino acid transporters were reported for their high expression and localization in the haustorial feeding structures. In regard to our study, it would be particularly pertinent to explore more precisely the balance between the different transporters in rust which are possibly involved in nitrogen acquisition.

## Acknowledgements

CL and SD are supported by the Labex Arbre (Programme Investissement d’Avenir, ANR-11-LABX-0002-01). CL thesis is funded by the Region Grand Est and the French National Research Agency (ANR-18-CE32-0001, Clonix2D project). SD and PF acknowledge the JGI Community Sequencing Program 2014 for the project 1416 *Further Studies in Poplar Rust Fungal Genomics*. The work (proposal: 10.46936/10.25585/60000832) conducted by the U.S. Department of Energy Joint Genome Institute (https://ror.org/04xm1d337), a DOE Office of Science User Facility, is supported by the Office of Science of the U.S. Department of Energy operated under Contract No. DE-AC02-05CH11231.

## Supplementary material

**Supplementary Figure S1.** Close-up on branches of clades 1 to 5 in the oligopeptide transporter (OPT) phylogenetic tree containing OPTs from selected Pucciniales species. OPTs from the family Melampsoraceae are depicted in pink whereas OPTs from other Pucciniales spp. are depicted in green.

**Supplementary Figure S2**. Expression profiling of *Melampsora larici-populina* transporter genes grouped by k-means across the life cycle. **A.** Genes are sorted by transporter classes and families according to the transporter classification database (TCDB). **B.** Genes are sorted by k-means (k= 10). Expression values scale from blue to red represents normalized expression levels from 0 (deep blue) to maximum value (deep red) at each stage. The Morpheus program parameters used for k-means were as follows: one minus Pearson’s correlation, cluster by rows, number of clusters 10 (1000 iterations). For each gene, the id shown on the right side of the profile indicates the Joint Genome Institute MycoCosm protein-ID number with the associated TCDB identifier and description. The 10 k-means are shown associated with a unique k-mean color on the right side of the profile. URE and UREg, dormant and germinating poplar rust urediniospores, respectively; T24 to T168, hours post-inoculation in a time course infection series on poplar leaves inoculated with urediniospores; TEL, telia; BSD, basidia; SPG and AEC, spermogonia and aecia formed on larch needles, respectively.

**Supplementary Table S1.** Published genomes in the Dikarya available in the JGI MycoCosm database on February 5^th^, 2022.

**Supplementary Table S2.** Transporter repertoires from 1051 Dikarya species available in the JGI MycoCosm database annotated according to the Transporter Classification Database (TCDB) on February 5^th^, 2022.

**Supplementary Table S3.** Expression values of transporter genes along the life cycle of the poplar rust fungus *Melampsora larici-populina* (extracted from Guerillot et al. 2021). URE and UREg, dormant and germinating poplar rust urediniospores, respectively; T24 to T168, hours post-inoculation in a time course infection series on poplar leaves inoculated with urediniospores; TEL, telia; BSD, basidia; SPG and AEC, spermogonia and aecia formed on larch needles, respectively.

**Supplementary Table S4.** Boxplot and statistical analyses for total genes, total transporter repertoires, and selected Transporter Classification Database (TCDB) classes and families.

**Supplementary Table S5.** Annotation of the *Melampsora allii-populina* oligopeptide transporter (OPT) gene family.

**Supplementary Table S6.** List of OPT genes retrieved from selected fungal species used in the phylogenetic analysis.

**Supplementary Table S7.** Table S7. List of the 361 OPT sequences used for the phylogenetic analysis with the OPT newick tree shown below sequences.

