## Supplementary figures and images for "A remarkable expansion of oligopeptide transporter genes in rust fungi (Pucciniales) suggests a specialization in nutrients acquisition for obligate biotrophy"

### Figure S1

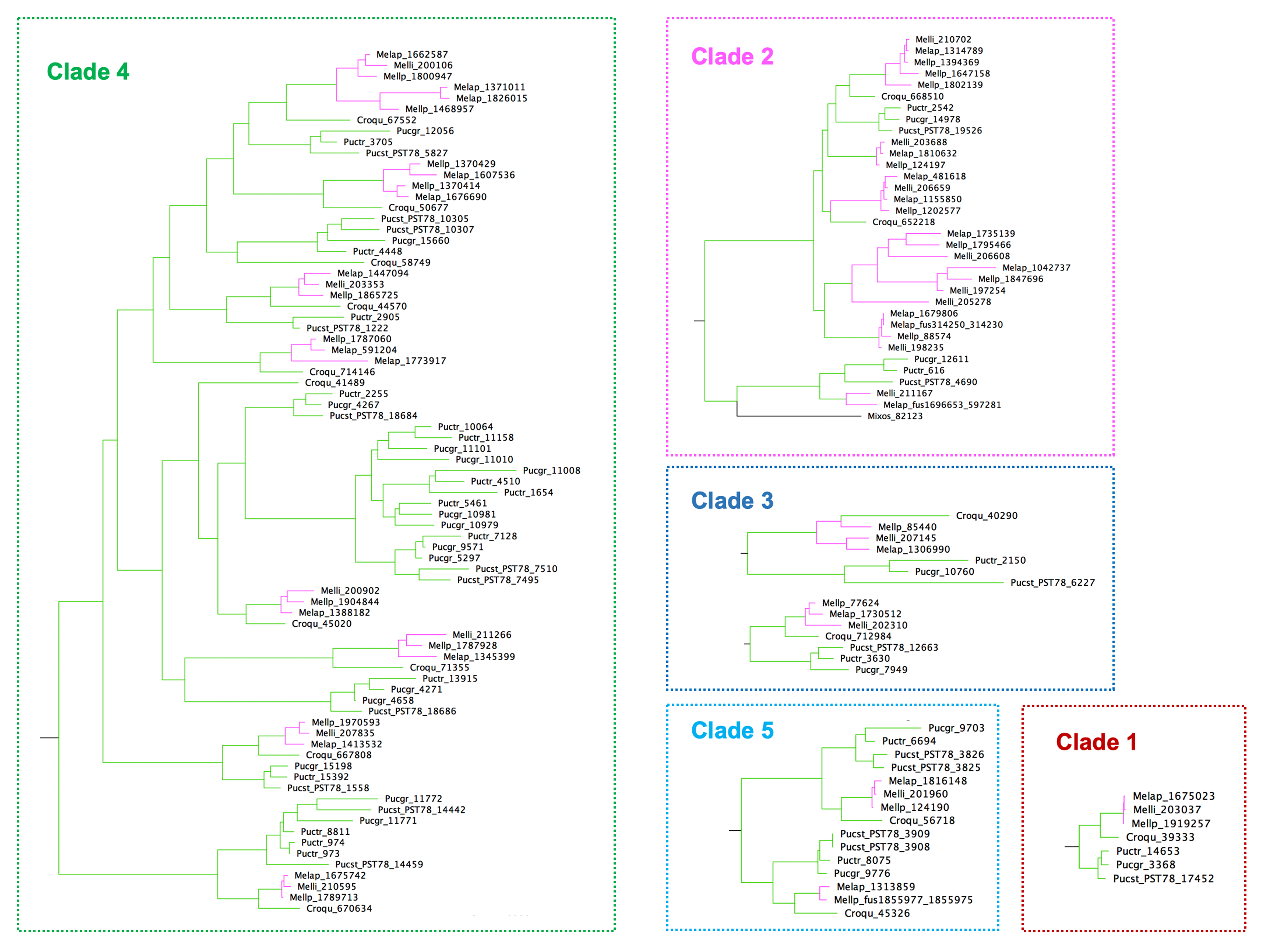

### Figure S2

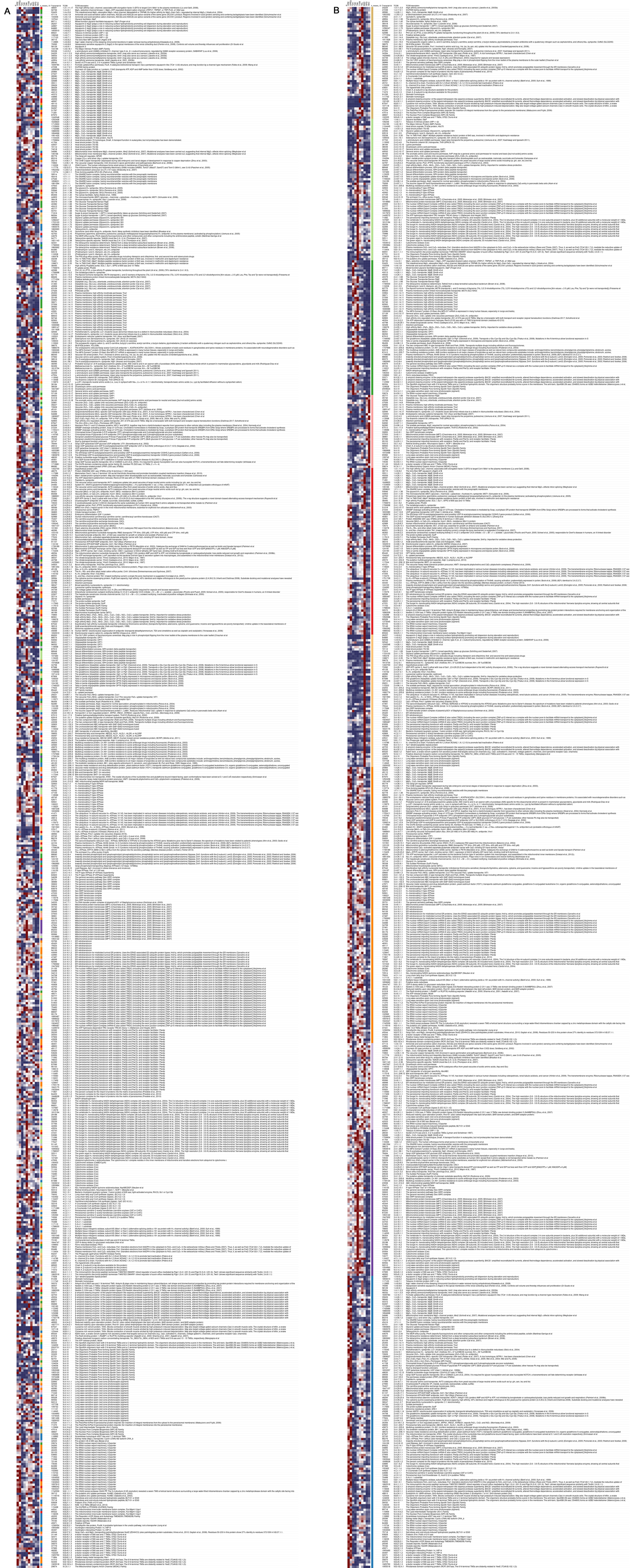
